# Constitutive DREADD signalling modulates oligodendrocyte precursor cell bioelectrical membrane properties and fate

**DOI:** 10.64898/2025.12.22.695316

**Authors:** Yasmine Kamen, Stavros Vagionitis, Omar de Faria, Ragnhildur Thóra Káradóttir

## Abstract

Oligodendrocyte precursor cells (OPCs) are small cells in the central nervous system that proliferate and differentiate into myelinating oligodendrocytes throughout life, allowing for myelin plasticity and repair. G protein-coupled neuromodulator receptors can regulate OPC fate, but the role of individual G protein families is unclear. Here, we use chemogenetics to directly investigate the role of Gq and Gi proteins in OPC fate. We find that expressing the DREADDs (Designer Receptors Exclusively Activated by Designer Drugs) hM3Dq or hM4Di in OPCs without activating them with designer agonists induces constitutive G protein signalling, which alters bioelectrical membrane properties such as voltage-gated ion channels and glutamate receptors in OPCs. Further, we find that hM3Dq or hM4Di expression alone modulates OPC fate, increasing or decreasing differentiation, respectively, suggesting that directly targeting G protein signalling can be used to bidirectionally regulate differentiation. Importantly, our data raise a note of caution regarding the use of DREADDs in both excitable and non-excitable small cells.

**SIGNIFICANCE STATEMENT:** Designer Receptors Exclusively Activated by Designer Drugs (DREADDs) are widely used to manipulate neuronal excitability. Their use in other neural cell types is rapidly increasing; however, DREADDs have not been thoroughly characterized in non-neuronal cells. Here, we show that when expressed in small excitable cells like oligodendrocyte precursor cells, DREADDs are constitutively active, unlike in excitatory neurons. This constitutive DREADD signalling alters cell fate and bioelectrical membrane properties like voltage-gated ion channels and glutamate receptors. Our data highlight that constitutive DREADD activity represents a significant confound that needs to be considered and controlled for when expressing DREADDs in small cells, and demonstrate that G protein signalling is a potent regulator of OPC fate, with a potential for therapeutic implications.

## INTRODUCTION

Lifelong myelination contributes to normal cognitive function and neural plasticity, and loss of myelin has been linked to cognitive decline and neurodegenerative disorders (de Faria et al., 2021). Oligodendrocyte precursor cells (OPCs) proliferate and differentiate into myelinating oligodendrocytes throughout life, underlying lifelong myelination (Young et al., 2013). Amongst other cues, neuromodulator signalling through G protein-coupled receptors (GPCRs) can regulate OPC proliferation and differentiation (Knapp and Hauser, 1996; Bongarzone et al., 1998; Ghiani et al., 1999; Ragheb et al., 2001; Stevens et al., 2002; Agresti et al., 2005; Luyt et al., 2007; Gomez et al., 2010, 2011, 2015; Niu et al., 2010; De Angelis et al., 2012; Coppi et al., 2013, 2020; Deshmukh et al., 2013; Mei et al., 2014, 2016a, 2016b; Spampinato et al., 2014; Vestal-Laborde et al., 2014; Abiraman et al., 2015; Tomas-Roig et al., 2016; Du et al., 2016; Chen et al., 2017; Welliver et al., 2018; Serrano-Regal et al., 2020; Osso et al., 2021; Seidman et al., 2022; Fiore et al., 2023; Lu et al., 2023; Bao et al., 2024; van de Wetering et al., 2024). However, neuromodulators activate multiple receptors coupled to different G proteins and thus, the role of individual G protein families in modulating OPC fate remains unclear. We sought to investigate the role of Gq and Gi proteins in regulating OPC proliferation, differentiation, and sensitivity to neuronal activity, and used DREADDs (Designer Receptor Exclusively Activated by Designer Drugs) to directly activate Gq or Gi signalling in OPCs.

Recent studies utilising the hM3Dq DREADD in OPCs have yielded contradictory results, with activation either reducing OPC proliferation and promoting differentiation (Fiore et al., 2023; Maas et al., 2025) or the reverse (Cheli et al., 2025). While the reasons underlying this discrepancy are unclear, one possibility is undetected constitutive activity. hM3Dq is a modified M3 muscarinic GPCR, and overexpressing GPCRs can induce constitutive G protein activity (Newman-Tancredi et al., 2000). Although DREADDs are not constitutively active in excitatory neurons (Alexander et al., 2009; Zhu et al., 2014), previous generations of designer receptors were constitutively active when expressed in smaller non-neuronal cells such as astrocytes (Sweger et al., 2007), cardiomyocytes (Redfern et al., 2000), or osteoblasts (Peng et al., 2008). Thus, it is conceivable that DREADDs could be constitutively active in OPCs. To investigate this, we expressed either the hM3Dq or hM4Di DREADDs in OPCs to drive Gq or Gi protein signalling, respectively, and tested whether these receptors altered OPC bioelectrical membrane properties and fate in the absence of an agonist. We found that expressing hM3Dq or hM4Di modulated OPC electrophysiological properties, and that these DREADD-induced changes could be reversed by G protein blockers, indicating that DREADDs likely induce constitutive G protein signalling. Importantly, hM3Dq and hM4Di expression also altered OPC proliferation and differentiation, suggesting that directly targeting Gq or Gi protein signalling can be used to bidirectionally modulate OPC fate. Our results highlight the importance of controlling for DREADD expression when designing experiments and raise a note of caution for DREADD data interpretation.

## RESULTS

### Expressing hM3Dq or hM4Di in OPCs alters their ion channels and glutamate receptors in the absence of a designer agonist

GPCRs can be constitutively active, and overexpressing a receptor can increase the probability of constitutive G protein signalling (Newman-Tancredi et al., 2000; Redfern et al., 2000). As DREADDs are modified muscarinic receptors and muscarinic receptors may modulate ion channels and glutamate receptors in OPCs (Kamen et al., 2024), we reasoned that constitutive DREADD activity in OPCs could result in altered bioelectrical membrane properties. Thus, we first examined whether hM3Dq and hM4Di expression altered OPC passive membrane properties, voltage-gated ion channels, and glutamate receptors. To do so, we whole-cell patch-clamped OPCs in the cingulate, motor, and somatosensory cortices of acute brain slices prepared from PdgfraCreER^T2^:hM3Dq-mCitrine/NG2-DsRed, PdgfraCreER^T2^:hM4Di-mCitrine/NG2-DsRed and NG2-DsRed control mice in the week following tamoxifen-induced recombination (Fig. 1a,b). We found that, in the absence of any agonist, hM3Dq expression decreased peak outward voltage-gated K^+^ (K_pk_) current density in OPCs without altering other bioelectrical membrane properties such as membrane resistance (Rm), resting membrane potential (Vm), inward K^+^ conductance (conductance_-74 mV to −154 mV_), steady-state outward K^+^ (K_ss_) current density, or voltage-gated Na^+^ channel (Na_V_) current density (Fig. 1c-i). hM4Di expression had a stronger effect on bioelectrical membrane properties, with Rm decreased, Vm becoming more negative, inward K^+^ conductance increased, and K_pk_ and Na_V_ densities decreased (Fig. 1j-o). hM4Di also increased both N-methyl-D-aspartate receptor (NMDAR) and α-amino-3-hydroxy-5-methyl-4-isoxazolepropionic acid/kainate receptor (AMPAR/KAR) current densities, in contrast to hM3Dq which did not alter glutamate receptors (Fig. 1p-u). These data suggest that hM3Dq and hM4Di expression alters OPC bioelectrical membrane properties, and therefore their sensitivity to neuronal activity, even in the absence of an agonist.

**Figure 1.**
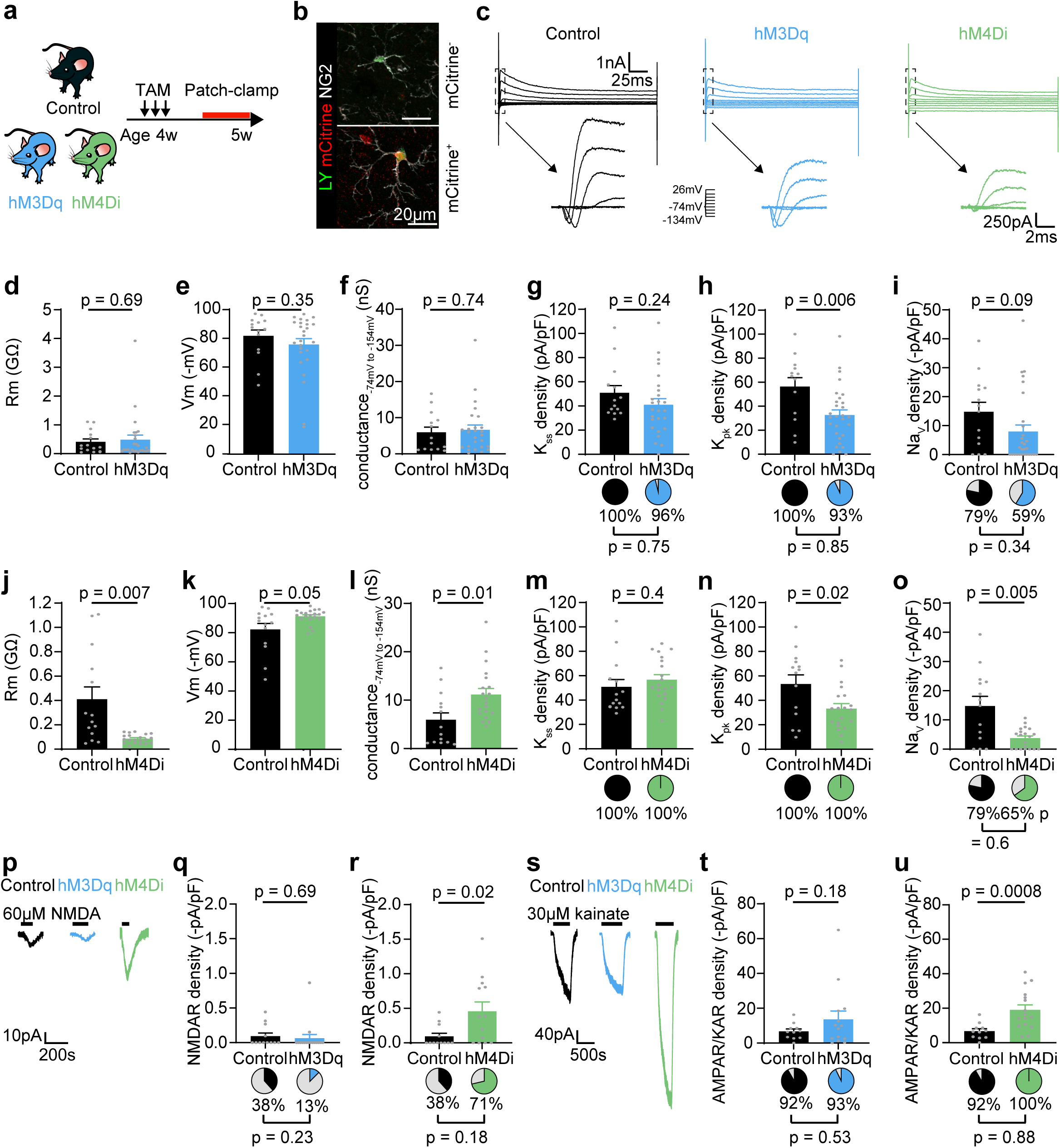
hM3Dq and hM4Di expression alter OPC bioelectrical membrane properties in the absence of an agonist. **a**, Experimental design. Control (NG2-DsRed), hM3Dq (PdgfraCreER^T2^:hM3Dq-mCitrine/NG2-DsRed or hM3Dq-mCitrine) and hM4Di (PdgfraCreER^T2^:hM4Di-mCitrine/NG2-DsRed) mice were dosed with tamoxifen daily for 3 days to induce DREADD expression. OPCs were whole-cell patch-clamped in the somatosensory, motor, and cingulate cortices after 3-11 days of expression. **b**, Example patch-clamped and Lucifer Yellow (LY)-filled control (mCitrine^−^) and DREADD-expressing (mCitrine^+^) NG2^+^ OPCs. **c**, Representative current traces in response to 20 mV voltage steps between −134 mV and +40 mV in control, hM3Dq and hM4Di OPCs. The arrows indicate the leak-subtracted currents during the first 7 ms of the voltage pulse. **d**-**f**, Quantification of membrane resistance (Rm, **d**), resting membrane potential (Vm, **e**), and inward K^+^ conductance (conductance_−74 mV to −154 mV_, **f**) in hM3Dq OPCs compared to control OPCs. **g**-**i**, Quantification of current density (bar graphs) and the proportion of OPCs with steady-state outward K^+^ (K_ss_) currents (**g**), peak outward voltage-gated K^+^ (K_pk_) currents (**h**), and voltage-gated Na^+^ channel (Na_V_) currents (**i**) in hM3Dq OPCs compared to control OPCs. **j**-**l**, Quantification of Rm (**j**), Vm (**k**), and conductance_−74 mV to −154 mV_ (**l**) in hM4Di OPCs compared to control OPCs. **m**-**o**, Quantification of K_ss_ current density and the proportion of cells with K_ss_ currents (**m**), K_pk_ current density and the proportion of cells with K_pk_ currents (**n**), and Na_V_ current density and the proportion of cells with Na_V_ (**o**) in hM4Di OPCs compared to control OPCs. **p**, Representative 60 µM NMDA-evoked currents in control, hM3Dq and hM4Di OPCs. **q**,**r**, Quantification of NMDAR current density (bar graphs) and the proportion of cells with NMDAR (pie charts) in control compared to hM3Dq (**q**) or hM4Di (**r**) OPCs. **s**, Representative 30 µM kainate-evoked currents in control, hM3Dq and hM4Di OPCs. **t**,**u**, Quantification of AMPAR/KAR current density (bar graphs) and the proportion of cells with AMPAR/KAR (pie charts) in control compared to hM3Dq (**t**) or hM4Di (**u**) OPCs. Data are show as mean±sem, with the grey dots indicating individual recorded cells. p-values were calculated by unpaired two-tailed t-test (bar graphs) or Yates’ corrected χ^2^ test (pie charts).

### DREADDs induce constitutive G protein signalling in OPCs

To investigate whether the DREADD-induced changes in OPC bioelectrical membrane properties resulted from constitutive Gq and Gi signalling in OPCs, we blocked Gq protein signalling in PdgfraCreER^T2^:hM3Dq-mCitrine/NG2-DsRed mice with the fast-acting blocker YM254890 (YM), and Gi protein signalling in PdgfraCreER^T2^:hM4Di-mCitrine/NG2-DsRed mice with pertussis toxin (Redfern et al., 2000) (PTX; Fig. 2a-c). We found that incubating acute brain slices prepared from PdgfraCreER^T2^:hM3Dq-mCitrine/NG2-DsRed mice in YM for one hour (Takasaki et al., 2004; Xiong et al., 2016) increased K_ss_ and inward K^+^ conductance in cortical OPCs but did not alter Rm, Vm, K_pk_ or Na_V_ (Fig 2d, f-k). This was surprising, as hM3Dq expression increased K_pk_ without altering K_ss_ or inward K^+^ conductance. Although YM is described as a selective Gq blocker, it has been suggested to block Gs and bind to G_βγ_ complexes released by GiPCRs (Peng et al., 2021), and might therefore act on endogenous GPCR signalling in OPCs. To validate that YM did not directly alter OPC bioelectrical membrane properties by altering endogenous GPCR signalling, we incubated acute brain slices prepared from control NG2-EYFP mice in YM for one hour, and found that YM had no effect on cortical OPC bioelectrical membrane properties (Fig. S1). Thus, YM only altered K_ss_ and inward K^+^ conductance when hM3Dq was expressed, suggesting that hM3Dq expression induces G protein constitutive signalling.

**Figure 2.**
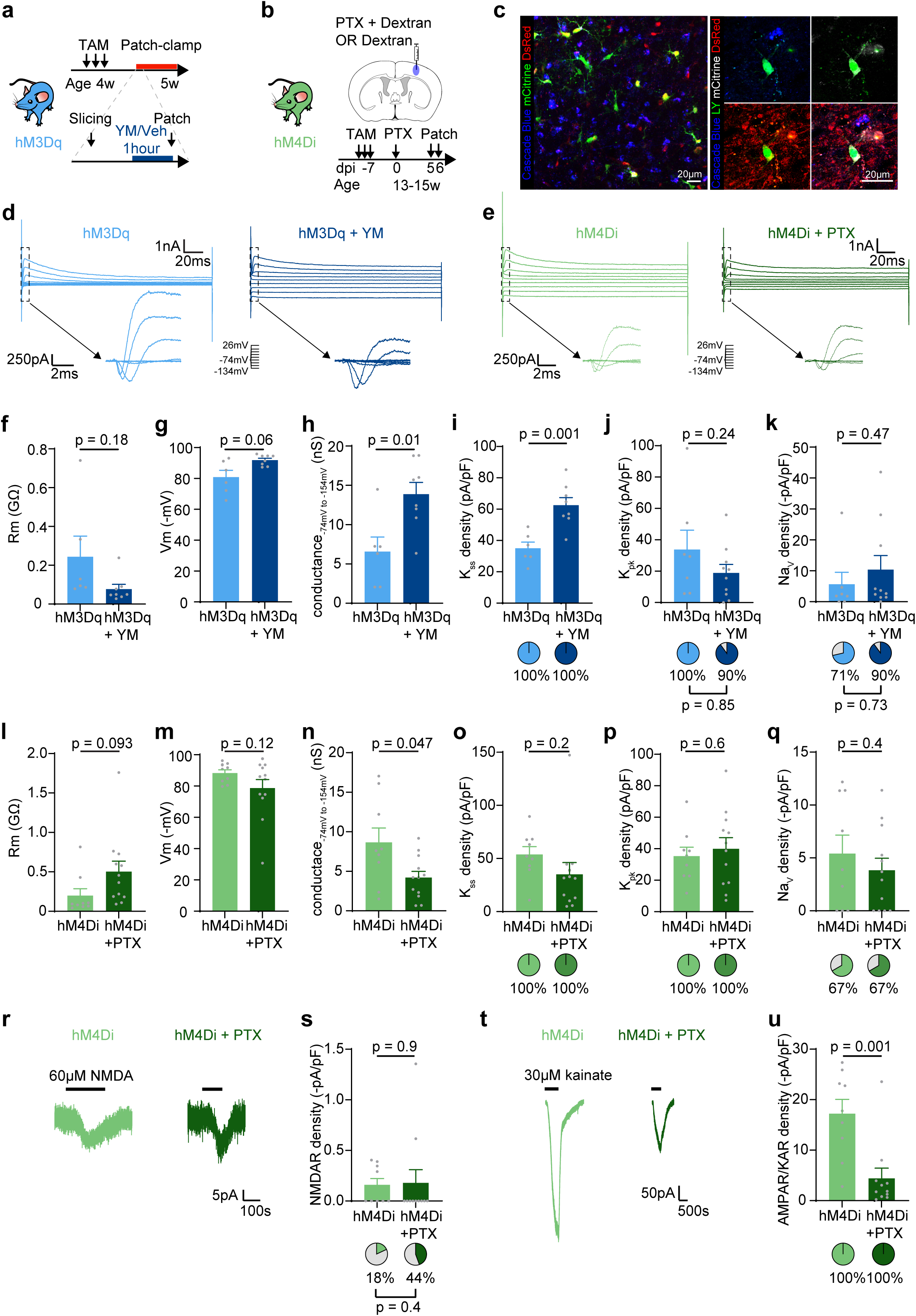
hM3Dq and hM4Di are constitutively active in OPCs. **a**, Experimental design to block Gq signalling. hM3Dq (PdgfraCreER^T2^:hM3Dq-mCitrine/NG2-DsRed) mice were dosed with tamoxifen daily for 3 days to induce DREADD expression. Acute brain slices were prepared after 3-7 days of expression, and incubated with 10 µM YM254890 (YM) or vehicle (0.1% DMSO) for one hour before cortical OPCs were whole-cell patch-clamped. **b**, Experimental design to block Gi signalling. hM4Di (PdgfraCreER^T2^:hM4Di-mCitrine/NG2-DsRed) mice were dosed with tamoxifen daily for 3 days to induce DREADD expression. 7 days later, 0.5 µg pertussis toxin (PTX) and Dextran Cascade Blue were injected stereotactically in the somatosensory cortex. OPCs surrounding Cascade Blue^+^ cells were whole-cell patch-clamped 5-6 days after the PTX injection. **c**, Example images with NG2-DsRed^+^ hM4Di-mCitrine^+^ OPCs near Cascade Blue^+^ cells (left) and a single patched and Lucifer Yellow (LY)-filled NG2-DsRed^+^ hM4Di-mCitrine^+^ OPC adjacent to a Cascade Blue^+^ cell (right). Scale bars 20 µm. **d**, Representative current traces in response to 20 mV voltage steps between −134 mV and +40 mV in hM3Dq and YM-treated OPCs. The arrows indicate the leak-subtracted currents during the first 7 ms of the voltage pulse. **e**, Representative current traces in response to 20 mV voltage steps between −134 mV and +40 mV in hM4Di and PTX-treated hM4Di OPCs. The arrows indicate the leak-subtracted currents during the first 7 ms of the voltage pulse. **f**-**k**, Quantification of membrane resistance (Rm, **f**), resting membrane potential (Vm, **g**), inward K^+^ conductance (conductance_−74 mV to −154 mV_, **h**), in YM-treated hM3Dq OPCs compared to hM3Dq OPCs. **o**-**q**, Quantification of current density (bar graphs) and the proportion of OPCs with steady-state outward K^+^ (K_ss_) currents (**i**), peak outward voltage-gated K^+^ (K_pk_) currents (**j**), and voltage-gated Na^+^ channel (Na_V_) currents (**k**) in YM-treated hM3Dq OPCs compared to hM3Dq OPCs. **l**-**q**, Quantification of Rm (**l**), Vm (**m**), conductance_−74 mV to −154 mV_ (**n**), K_ss_ current density and the proportion of cells with K_ss_ currents (**o**), K_pk_ current density and the proportion of cells with K_pk_ currents (**p**), and Na_V_ current density and the proportion of cells with Na_V_ (**q**) in PTX-treated hM4Di OPCs compared to hM4Di OPCs. **r**, Representative 60 µM NMDA-evoked currents in hM4Di and PTX-treated hM4Di OPCs. **s**, Quantification of NMDAR current density (bar graphs) and the proportion of cells with NMDAR (pie charts) in hM4Di compared to PTX-treated hM4Di OPCs. **t**, Representative 30 µM kainate-evoked currents in hM4Di and PTX-treated OPCs. **u**, Quantification of AMPAR/KAR current density (bar graphs) and the proportion of cells with AMPAR/KAR (pie charts) in hM4Di and PTX-treated hM4Di OPCs. Data are shown as mean±sem, with the grey dots indicating individual recorded cells. p-values were calculated by unpaired two-tailed t-test (bar graphs) or Yates’ corrected χ^2^ test (pie charts).

To block Gi protein in hM4Di-expressing OPCs, we injected PTX in the somatosensory cortex of PdgfraCreER^T2^:hM4Di-mCitrine/NG2-DsRed mice and whole-cell patch-clamped OPCs 5-6 days later (Funada et al., 1993; Lee et al., 2015) (Fig. 2b,c). Blocking Gi signalling with PTX reversed the effect of hM4Di expression on inward K^+^ conductance and AMPAR/KAR current density, but had no effect on other bioelectrical membrane properties, although we observed a trend in Rm and Vm (Fig 2e, l-u), perhaps due to involvement of the PTX-insensitive Gi family member Gz (Casey et al., 1990). Of note, both inward K^+^ conductance and AMPAR/KAR density were reduced to the same level as in control mice (inward K^+^ conductance: 5-week NG2-DsRed control: 5.94±1.44 nS, n=14, 13-15-week hM4Di+PTX: 4.20±0.79 nS, n=12, p=0.3; AMPAR/KAR density: 5-week NG2-DsRed control: 6.70±1.37 pA/pF, n=11, 13-15-week hM4Di+PTX: 4.42±2.03 pA/pF, n=11, p=0.4; unpaired two-tailed t-tests), suggesting that hM4Di-induced Gi constitutive signalling drives the increase in inward K^+^ conductance and AMPAR/KAR density in hM4Di OPCs.

### Expressing hM3Dq reduces OPC proliferation and increases differentiation

As bioelectrical membrane properties modulate OPC fate (Chen et al., 2018; Khawaja et al., 2021; Pivoňková et al., 2024), we next tested whether hM3Dq- and hM4Di-driven constitutive G protein signalling altered OPC proliferation and differentiation. We induced hM3Dq and hM4Di expression in 4-week-old mice, and perfused-fixed these mice 3 weeks later (Fig. 3a). We took advantage of sparse recombination in PdfraCreER^T2^:hM3Dq-mCitrine mice (8% of OPCs were recombined 3-7 days post-tamoxifen; 3% of OPCs and 10% of OLIG2^+^ cells were recombined 3 weeks post-tamoxifen), to compare the fate of recombined OPCs (mCitrine^+^) to non-recombined OPCs (mCitrine^−^) in the same environment, as well as in control PdgfraCreER^T2^ littermates. To assess proliferation, we injected the mice with EdU (5-Ethynyl-2’-deoxyuridine) 24 hours prior to perfusion-fixation (Fig. 3a). We found that hM3Dq expression significantly reduced EdU incorporation in cingulate cortex OPCs, indicating a decrease in proliferation (Fig. 3b,c). In addition, hM3Dq expression significantly increased OPC differentiation into CC1^+^ oligodendrocytes (Fig. 3d,e). Surprisingly, non-recombined OPCs in PdfraCreER^T2^:hM3Dq-mCitrine mice differentiated less than in control littermates, suggesting that excessive differentiation by hM3Dq-expressing OPCs blocked differentiation of non-recombined cells. This resulted in an overall decrease in differentiation and myelin basic protein (MBP) positive area in the cingulate cortex of PdfraCreER^T2^:hM3Dq-mCitrine mice compared to control PdfraCreER^T2^ littermates (differentiation (% CC1^+^OLIG2^+^/OLIG2^+^): control mice: 56.92±1.46%, n=3, hM3Dq mice: 42.59±4.09%, n=4, p=0.03; MBP^+^ area: control mice: 46.43±1.71%, n=3, hM3Dq mice: 40.83±1.14%, n=4, p=0.04; unpaired two-tailed t-tests). As hM3Dq expression increased differentiation but did not result in an overall increase in myelination, we examined whether cell death was increased following hM3Dq expression. We only detected a total of 3 cleaved caspase-3^+^OLIG2^+^ cells across 8 animals and found no cleaved caspase-3^+^ hM3Dq-expressing cells (hM3Dq, n=4 mice, control, n=4 mice; 8 images per mouse), suggesting that hM3Dq expression does not induce caspase-3 mediated cell death. Taken together, these data indicate that hM3Dq expression alters OPC fate in the absence of an agonist.

**Figure 3.**
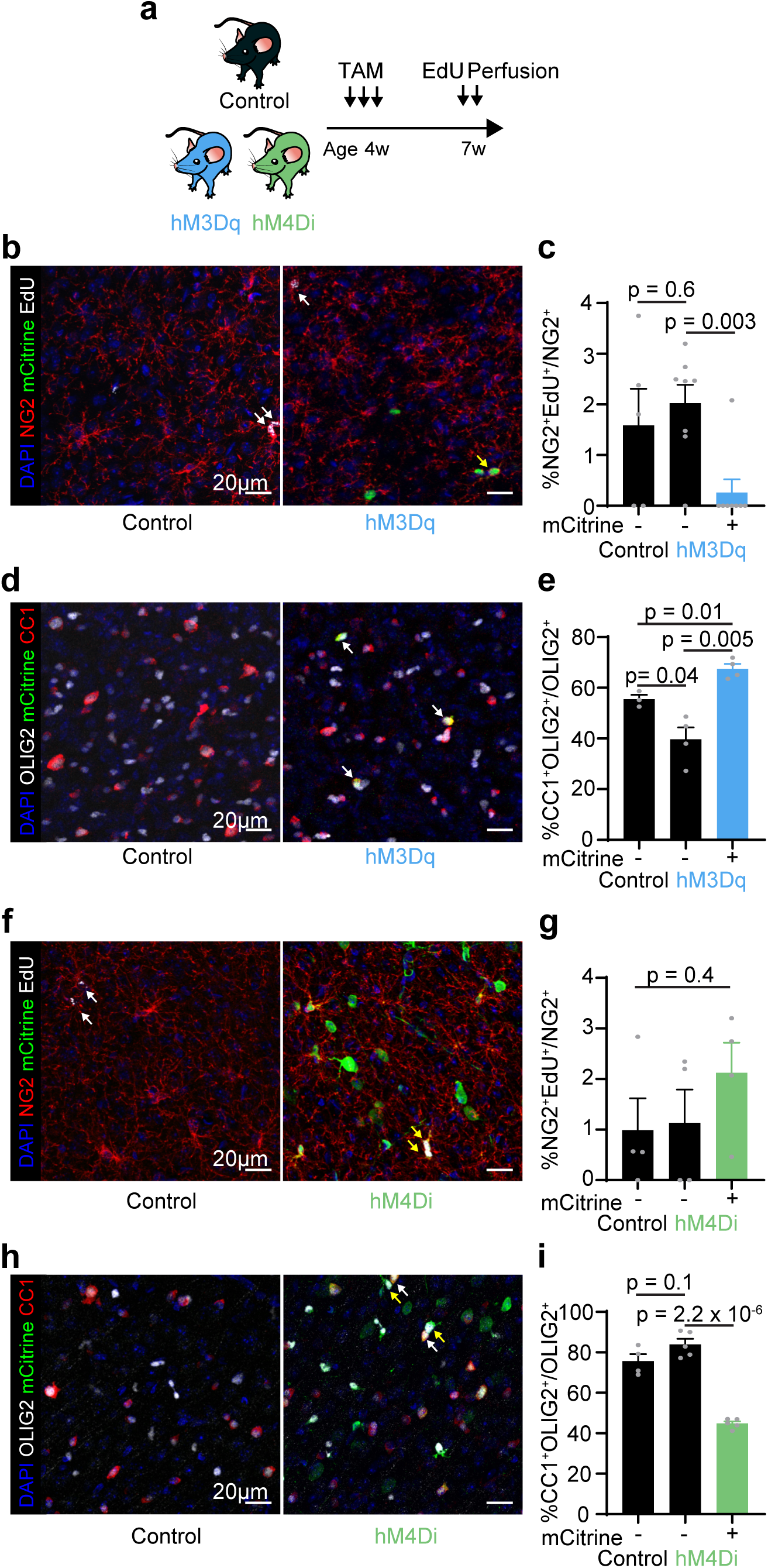
hM3Dq and hM4Di constitutive signalling modulates OPC fate. **a**, Experimental design. hM3Dq (PdgfraCreER^T2^:hM3Dq-mCitrine), hM4Di (PdgfraCreER^T2^:hM4Di-mCitrine) and littermate control (PdgfraCreER^T2^) mice were dosed with tamoxifen daily for 3 days to induce DREADD expression. Mice were perfused-fixed for immunohistochemistry 3 weeks later, with EdU administration 24 hours prior to perfusion. **b**, Representative images of NG2^+^EdU^+^ cells in the cingulate cortex of control and hM3Dq mice. The white arrows indicate NG2^+^EdU^+^mCitrine^−^ cells in control and hM3Dq mice. The yellow arrow indicates an NG2^+^EdU^−^mCitrine^+^ cell in an hM3Dq mouse. Scale bars 20 µm. **c**, Quantification of proliferation in mCitrine^+^ OPCs compared to mCitrine^−^ OPCs in control or hM3Dq mice (p=0.01, one-way ANOVA). **d**, Representative images of CC1^+^OLIG2^+^ cells in the cingulate cortex of control and hM3Dq mice. The white arrows indicate CC1^+^OLIG2^+^mCitrine^+^ cells in an hM3Dq mouse. Scale bars 20 µm. **e**, Quantification of differentiation in mCitrine^+^ cells compared to mCitrine^−^ cells in control or hM3Dq mice (p=9×10^−4^, one-way ANOVA). **f**, Representative images of NG2^+^EdU^+^ cells in the cingulate cortex of control and hM4Di mice. The white arrows indicate NG2^+^EdU^+^mCitrine^−^ cells in a control mouse. The yellow arrows indicate NG2^+^EdU^+^mCitrine^+^ cells in an hM4Di mouse. Scale bars 20 µm. **g**, Quantification of proliferation in mCitrine^+^ OPCs compared to mCitrine^−^OPCs in control or hM4Di mice (p=0.4, one-way ANOVA). **h**, Representative images of CC1^+^OLIG2^+^ cells in the cingulate cortex of control and hM4Di mice. The white arrows indicate example CC1^+^OLIG2^+^mCitrine^+^ cells in an hM4Di mouse. The yellow arrows indicate example CC1^+^OLIG2^+^mCitrine^+^ cells in an hM4Di mouse. Scale bars 20 µm. **i**, Quantification of differentiation in mCitrine^+^ cells compared to mCitrine^−^ cells in control or hM4Di mice (p=4.9×10^−7^, one-way ANOVA). Data are shown as mean ± sem. Grey data points indicate individual animals. p-values on the bar graphs are from Holm-Bonferroni post-hoc tests following a statistically significant one-way ANOVA (c, e, i) or from a one-way ANOVA (g).

### Expressing hM4Di reduces differentiation

As with hM3Dq, we found that expressing hM4Di in OPCs modulated their fate. Using a similar approach with PdfraCreER^T2^:hM4Di-mCitrine mice (51% of OPCs were recombined 3-7 days post-tamoxifen; 74% of OPCs and 29% of OLIG2^+^ cells were recombined 3 weeks post-tamoxifen), we found that hM4Di expression did not alter proliferation in the cingulate cortex (Fig. 3f,g), but significantly decreased differentiation into CC1^+^ oligodendrocytes (Fig. 3h,i). However, this did not have a detectable effect on overall CC1^+^ oligodendrocyte density or MBP density in the cingulate cortex of PdfraCreER^T2^:hM4Di-mCitrine mice compared to control PdfraCreER^T2^ littermates (differentiation (% CC1^+^OLIG2^+^/OLIG2^+^): control mice: 75.69±3.45%, n=4, hM4Di mice: 71.03±2.42%, n=5, p=0.3; MBP^+^ area: control mice: 17.90±2.24%, n=4, hM4Di mice: 16.44±1.63%, n=5, p=0.6; unpaired two-tailed t-test), perhaps due to compensation by non-recombined cells or to differentiation prior to tamoxifen administration. Of note, we detected recombination in this line in a small proportion of neurons (10% of NeuN^+^ neurons expressed hM4Di-mCitrine) resulting from spontaneous recombination of the hM4Di-mCitrine transgene, as this was detected without tamoxifen administration but was not detected in PdfraCreER^T2^:hM3Dq-mCitrine mice (Fig. S2). This small proportion of recombined neurons is unlikely to have modulated OPC fate as the proliferation and differentiation of non-recombined OPCs in PdfraCreER^T2^:hM4Di-mCitrine mice did not differ from that of OPCs in control mice (Fig. 3h,i). This is consistent with DREADD expression not inducing constitutive G protein activity in neurons (Alexander et al., 2009; Zhu et al., 2014), and suggests that hM4Di expression in OPCs directly reduces their differentiation.

## DISCUSSION

We sought to determine whether directly activating Gq or Gi protein signalling in OPCs could alter their fate and sensitivity to neuronal activity. By using the hM3Dq and hM4Di DREADDs to do so, we identified that DREADD expression alone induces constitutive G protein signalling in OPCs and modulates OPC passive membrane properties, voltage-gated ion channels, and glutamate receptors, resulting in altered proliferation and differentiation dynamics. This raises an important technical concern with the use of DREADDs, but also suggests that directly activating Gq or Gi protein signalling in OPCs can modulate their sensitivity to neuronal activity, as well as their proliferation and differentiation. OPCs exhibit regional and temporal heterogeneity in their electrophysiological properties (Chittajallu et al., 2004; Karadottir et al., 2008; Spitzer et al., 2019; Pivoňková et al., 2024), response to growth factors (Hill et al., 2013), and proliferation and differentiation potential (Viganò et al., 2013; Young et al., 2013), which could represent different cell types (Marisca et al., 2020) or plastic functional cell states (Kamen et al., 2022). Our data support OPCs existing in multiple functional states, as we found that hM3Dq and hM4Di expression reversibly shifted OPCs between different ion channel profiles and modulated cell fate. We have previously suggested that pro-remyelination compounds administered systemically act by modulating OPC states (Kamen et al., 2024), and here, we show that directly manipulating how OPCs sense their environment alters their fate. Overall, our data indicate that directly targeting G protein activity in OPCs may be a therapeutic target to bidirectionally regulate their fate.

Our data raise a note of caution for the use of DREADDs and the interpretation of results. For instance, we found that hM3Dq expression alone increased differentiation at the expense of proliferation, a phenotype that has been reported when comparing clozapine-N-oxide treated hM3Dq^+^ OPCs to clozapine-N-oxide treated hM3Dq^−^ OPCs (Fiore et al., 2023; Maas et al., 2025). In contrast, comparing vehicle treated hM3Dq^+^ OPCs to clozapine-N-oxide treated hM3Dq^+^ OPCs suggests that Gq activation promotes proliferation at the expense of differentiation (Cheli et al., 2025). Although other aspects of the experimental design may contribute to the differences between these reports, our data highlight the importance of accounting for designer receptor expression when working with DREADDs.

We found that blocking G protein signalling with YM or PTX partially reversed DREADD-induced electrophysiological changes, suggesting that DREADD expression induces constitutive G protein signalling. OPCs may be particularly prone to constitutive G protein signalling, as numerous drug screening studies report GPCR antagonists altering oligodendrocyte differentiation without exogenous agonists: blocking the Gi-coupled D2/3 receptors or the Gq-coupled M1 receptor in isolated OPC cultures can alter OPC fate in the absence of agonist (Niu et al., 2010; Deshmukh et al., 2013; Mei et al., 2014), and Gi-coupled H3 receptor constitutive signalling decreases OPC differentiation (Chen et al., 2017). Previous generations of designer receptors induced constitutive activity in non-neuronal cells (Redfern et al., 2000; Sweger et al., 2007; Peng et al., 2008) and it has been suggested that high viral titers or strong promoters might induce DREADD constitutive activity (Roth, 2016). Here, we used readily available Cre-dependent mouse lines (Zhu et al., 2016) to express hM3Dq and hM4Di; in these lines, hM3Dq and hM4Di expression is driven by a generic CAG promoter. However, OPCs are small cells, and thus, expression levels that do not induce constitutive activity in excitatory neurons may be sufficient to induce constitutive activity in smaller cells, potentially also including microglia, astrocytes, or interneurons alongside OPCs. Using mouse lines with cell type-specific or weaker promoters could potentially help prevent constitutive DREADD activity (Roth, 2016). Our data indicate that controlling for any potential constitutive activity is critical when expressing DREADDs in small cells, especially as hM3Dq or hM4Di altered OPCś passive bioelectrical membrane properties, voltage-gated ion channels and glutamate receptors, which are hallmarks of most neural cells.

## ACKNOWLEDGEMENTS

We thank Prof. William Richardson for the PdgfraCreER^T2^ mice and Prof. Jacqueline Trotter for the NG2-EYFP mice. We acknowledge the support of the Cambridge Stem Cell Institute core facility staff and University Biomedical Services. This project has received funding from: the European Research Council (ERC) under the European Union’s Horizon 2020 research and innovation programme (grant agreement No 771411; R.T.K., Y.K.); the Wellcome Trust, a studentship (102160/Z/13/Z; Y.K.); the Fonds de recherche du Québec-Santé, a scholarship (Y.K.); the Cambridge Commonwealth European & International Trust, a scholarship (Y.K.); MS Society Centre Excellence grant, Cambridge Myelin Repair Centre grant (132; R.T.K., O.d.F., S.V.); and the Lister Institute, a Research Prize (R.T.K.). This research was supported by the Wellcome Trust (203151/Z/16/Z, 203151/A/16/Z) and the UKRI Medical Research Council (MC_PC_17230) Cambridge Stem Cell Institute core funding. For the purpose of open access, the author has applied a CC BY public copyright licence to any Author Accepted Manuscript version arising from this submission.

## AUTHOR CONTRIBUTIONS

Conceptualization, Y.K., R.T.K.; Investigation, Y.K., O.d.F., S.V.; Data analysis: Y.K., R.T.K; Writing, Y.K., R.T.K., and all authors commented on the manuscript; Funding Acquisition, Resources and Supervision, R.T.K.

## METHODS

### Animals

Experiments were performed in accordance with EU guidelines for the care and use of laboratory animals, and with the guidelines of the UK Animals (Scientific Procedures) Act 1986 and subsequent amendments. Use of animals in this project was approved by the Animal Welfare and Ethical Review Body for the University of Cambridge and carried out under the terms of UK Home Office Licenses and in line with ARRIVE guidelines. All mice were maintained under a 12h light:12h dark cycle with food and water supplied *ad libitum*. The following mice were obtained from Jackson Laboratory: hM3Dq-mCitrine (B6N;129-Tg(CAG-CHRM3*,-mCitrine)1Ute/J; JAX 026220) (Zhu et al., 2016) and hM4Di-mCitrine (B6.129-Gt(ROSA)26Sor^tm1(CAG-CHRM4*,-mCitrine)Ute^/J; JAX 026219) (Zhu et al., 2016), and bred in house with PdgfraCreER^T2^ mice (Tg(Pdgfra-cre/ERT2)1Wdr) (Rivers et al., 2008), a gift from Prof. William Richardson (University College London, United Kingdom). For patch-clamp experiments, these mice were further crossed with NG2-DsRed mice (Tg(Cspg4-DsRed.T1)1Akik/J; JAX 008241) (Zhu et al., 2008). NG2-EYFP mice (Karram et al., 2008), a gift from Prof. Jacqueline Trotter, were used to test the effect of YM254890 on control OPCs (Figure S1).

### Induction of Cre-mediated recombination

For the experiments shown in Figure 1, three- to four-week-old PdgfraCreER^T2^:hM3Dq-mCitrine/NG2-DsRed and PdgfraCreER^T2^:hM4Di-mCitrine/NG2-DsRed were given daily doses of 150 mg/kg tamoxifen by oral gavage for three consecutive days. Age-matched control NG2-DsRed mice received the same tamoxifen dosing. Patch-clamp experiments took place three to seven days after the last day of tamoxifen dose, in P26 to P35 mice. Five of 25 patched hM3Dq cells were from hM3Dq-mCitrine mice that were injected intraperitoneally with 20 mg/kg tamoxifen daily between P7 and P9 and patch-clamp experiments were performed 10 and 11 days after the last tamoxifen injection in P19 and P20 mice. These cells were pooled with the PdgfraCreER^T2^:hM3Dq-mCitrine/NG2-DsRed cells. For the experiments shown in Figure 2, three- to four-week-old PdgfraCreER^T2^:hM3Dq-mCitrine/NG2-DsRed mice were given daily doses of 150 mg/kg tamoxifen by oral gavage for three consecutive days and patch-clamp experiments were performed four to six days following the last tamoxifen dose in P27 to P29 mice. 12- to 13-week-old PdgfraCreER^T2^:hM4Di-mCitrine/NG2-DsRed were given daily doses of 150 mg/kg tamoxifen by oral gavage for three consecutive days. Intracranial injections were performed seven days after the final tamoxifen dose, and patch-clamp recordings were performed five and six days following surgery, in P97 to P113 mice. For the experiments shown in Figure 3, three- to four-week-old PdgfraCreER^T2^:hM3Dq-mCitrine mice, PdgfraCreER^T2^:hM4Di-mCitrine mice, and PdgfraCreER^T2^ littermate control mice were given daily doses of 150 mg/kg tamoxifen by oral gavage or 100 mg/kg tamoxifen by intraperitoneal injection for three consecutive days.

### Electrophysiology

225µm-thick coronal slices were prepared from PdgfraCreER^T2^:hM3Dq-mCitrine, PdgfraCreER^T2^:hM3Dq-mCitrine/NG2-DsRed, PdgfraCreER^T2^:hM4Di-mCitrine/NG2-DsRed, and NG2-DsRed mice between postnatal day 19 (P19) and P113, as specified in the figures, in ice-cold (∼1°C) oxygenated (5% CO_2_ / 95% O_2_) bicarbonate-buffered aCSF containing, in mM: 124 NaCl, 26 NaHCO_3_, 1 NaH_2_PO_4_, 2.5 KCl, 2 MgCl_2_, 2.5 CaCl_2_, 10 glucose, pH 7.4, 330 mOsm. 1 mM kynurenic acid was added to block glutamate receptors that might be activated during dissection (Kamen and Káradóttir, 2021). Slices were allowed to rest for at least one hour before starting recordings. All experiments were performed in whole-cell voltage-clamp mode, with holding potential −74 mV. Cells were recorded in the cingulate, motor, and somatosensory cortices. Pipette resistance was between 4.7-6.5 MΩ and mean uncompensated series resistance was 25 MΩ. Recordings were performed in HEPES-buffered aCSF containing, in mM: 144 NaCl, 10 HEPES, 1 NaH_2_PO_4_, 2.5 KCl, 2.5 CaCl_2_ and 10 glucose, with pH adjusted to 7.2-7.4 with 1 M NaOH, and osmolarity 315 mOsm. Mg^2+^ was omitted to record NMDA-evoked responses. The internal solution contained, in mM: 130 K-gluconate, 4 NaCl, 0.5 CaCl_2_, 10 HEPES, 10 BAPTA, 4 Mg_x_ATP, 0.5 Na_x_GTP, and 2 K-Lucifer Yellow, pH adjusted to 7.2-7.4 with 2 M KOH, with osmolarity between 290 and 300 mOsm. All recordings took place at room temperature, and the recording solution was continuously oxygenated with 100% O_2_. Inclusion criteria were based on series resistance, leak current being lower than 400 pA and a stable baseline. An Axopatch 200 (Molecular Devices) was used for voltage-clamp data acquisition. Voltage step data were sampled at 50 kHz and filtered at 10 kHz and drug application data were sampled at 1 kHz using pClamp 10.7 or pClamp 11 (Molecular Devices). Cells were recorded in cortical layers 2-6. During recordings, cells were filled with Lucifer Yellow. Location and cell identity were confirmed by post-hoc immunohistochemistry against mCitrine and NG2. In 41/41 cases, imaged cells were positive for NG2 and/or NG2-DsRed, and in 33/33 cases imaged DREADD cells were also positive for mCitrine.

100 µM glycine, an NMDA co-agonist, was included in the recording solution. 5 µM strychnine, a glycine receptor antagonist, was also included to avoid activation of glycine receptors. 60 µM NMDA was used to activate NMDAR, and 30 µM kainate was used to activate both AMPAR and KAR. 500 µM BaCl_2_ was added to the recording solution after determination of passive membrane properties to block inward K^+^ conductance (Kamen and Káradóttir, 2021).

Series resistance, membrane resistance, and membrane capacitance were calculated as previously described with a custom MATLAB script (Agathou and Káradóttir, 2019; Spitzer et al., 2019; Kamen and Káradóttir, 2021). Peak voltage-gated Na^+^ channel currents and peak outward voltage-gated K^+^ channel currents were measured on leak-subtracted currents (Kamen and Káradóttir, 2021) during the first 7 ms of the voltage pulse, and steady-state outward K^+^ channel currents were measured on unsubtracted currents 170 ms to 200 ms after the onset of the voltage pulse; all subtractions and measurements were performed with custom MATLAB scripts (Spitzer et al., 2019). Membrane potential and inward K^+^ conductance (calculated from the line of best fit between −154 mV and the equilibrium potential) were determined manually in Excel (Pivoňková et al., 2024). Glutamate receptor currents were measured manually in pClamp at the peak of the agonist response, with a baseline drift correction (Agathou and Káradóttir, 2019; Kamen and Káradóttir, 2021).

### YM254980 treatment

To block Gq protein signalling, acute brain slices were incubated in 10 µM YM254890 and 0.1% DMSO in standard slicing solution for one hour (Takasaki et al., 2004; Xiong et al., 2016). Control slices were incubated in 0.1% DMSO for one hour.

### PTX treatment

PdgfraCreER^T2^:hM4Di-mCitrine/NG2-DsRed mice aged P94 or P106 were anesthetized with 2% Isoflurane and injected stereotaxically into the somatosensory cortex (antero-posterior at bregma, mediolateral 2.25 mm, dorso-ventral 1.4 mm, 200 nl/min) with 0.5 µg PTX (Funada et al., 1993; Lee et al., 2015) (Tocris, 3097) and 0.025 µg Dextran Cascade Blue (ThermoFisher Scientific, D1976) to label the injection site or Dextran Cascade Blue only (control) into the somatosensory cortex. Each animal received a single injection. Meloxicam was given pre-operatively and for three days post injection for pain relief. Electrophysiological recordings were performed five and six days after the injection.

### EdU administration to mice

PdgfraCreER^T2^:hM3Dq-mCitrine, PdgfraCreER^T2^:hM4Di-mCitrine, and PdgfraCreER^T2^ littermate controls were injected intraperitoneally with 50 mg/kg EdU (Mitew et al., 2018) (Life Technologies, E10187; dissolved in 0.9% saline) 24 hours prior to perfusion-fixation.

### Immunohistochemistry and imaging

Seven-week-old mice were anesthetized with 5% isoflurane and injected intraperitoneally with 20 mg pentobarbital sodium before perfusion-fixation with ice-cold PBS followed by ice-cold 4% PFA. Dissected brains were postfixed for one hour at room temperature before being washed in PBS. 100 µm-thick slices were cut on a vibrating microtome. Alternatively, acute brain slices were incubated in 4% PFA for one hour at room temperature, before being washed in PBS. For antibody labelling, free-floating slices were incubated in 10% goat or donkey serum and 0.5% Triton-X 100 in PBS for 4-5 hours at room temperature, on a rotating shaker. Slices were incubated with primary antibodies in PBS overnight at room temperature. Primary antibodies were as follows: chicken anti-GFP, 1:500 or 1:1000 (Abcam, ab13970); rabbit anti-OLIG2, 1:300 (EMD Millipore, AB9610); goat anti-OLIG2, 1:300 (R&D Systems, AF2418); rat anti-MBP, 1:100 (AbD Serotec, MCA409S); rabbit anti-NG2, 1:300 (Millipore, AB5320); mouse anti-APC, clone CC1, 1:300 (Millipore, MABC200 or OP80); mouse anti-NeuN 1:300 (EMD Millipore, MAB377). Following three 30 minutes washes in PBS, the slices were incubated in secondary antibodies in PBS at a 1:1000 concentration overnight at 4°C or for 5 hours at room temperature, on a rotating shaker. Secondary antibodies were as follows: goat anti-chicken IgY Alexa Fluor 448 (Abcam, ab150169), goat anti-chicken IgY Alexa Fluor 568 (Invitrogen, A-11041), goat anti-chicken IgY Alexa Fluor 647 (Invitrogen, A-21449), goat anti-rabbit IgG Alexa Fluor 488 (Invitrogen, A-11008), goat anti-rabbit IgG Alexa Fluor 555 (Invitrogen, A-21429), goat anti-rabbit IgG Alexa Fluor 568 (Invitrogen, A-11036), goat anti-rabbit IgG Alexa Fluor 647 (Invitrogen, A-21245), goat anti-rat IgG Alexa Fluor 488 (Invitrogen A-11006), goat anti-mouse IgG Alexa Fluor 633 (Invitrogen, A-21052), goat anti-mouse IgG Alexa Fluor 647 (Invitrogen, A-21236), donkey anti-goat IgG Alexa Fluor 488 (Invitrogen, A-11055), donkey anti-rabbit IgG Alexa Fluor 555 (Abcam, ab150074), and donkey anti-chicken IgY Alexa Fluor 647 (Stratech, 703-605-155-JIR). After two washes in PBS, slices were incubated with 1 ng/ml DAPI for 20 minutes, and following a final PBS wash, mounted on glass slides.

To visualise EdU-labelled cells, we used a Click-iT Cell Reaction Buffer Kit (Invitrogen, C10269). Following antibody staining, and immediately after DAPI incubation, slices were washed in 2% BSA for 10 minutes, then incubated for 30 minutes with the Click It reaction mix (prepared according to the manufacturer’s instructions; we included an Alexa Fluor 555-conjugated Azide (Invitrogen, A20012)). Following this, slices were washed for 10 minutes with 2% BSA, and then for 30 minutes with PBS before mounting them on glass slides.

Antigen retrieval was performed for cleaved caspase-3 staining (rabbit anti cleaved caspase-3, 1:750, Cell Signaling Technology, 9661), with slices incubated in pre-heated (98 °C) 10 mM TRIS, 1 mM EDTA, and 0.05% TWEEN in PBS buffer at pH 9 for three minutes and allowed to cool down at room temperature in antigen retrieval solution before overnight incubation at 4°C on a rotating shaker with primary antibodies in 1% bovine serum albumin (BSA) and 0.3% Triton X 100. Following three PBS washes, slices were incubated with secondary antibodies in 1% BSA and 0.3% Triton X 100 for one hour at room temperature on a rotating shaker before a further two PBS washes, incubation with DAPI, and mounting.

Samples were imaged on a Leica TCS SP5 microscope or a Leica TCS SP8 microscope. Laser intensity, voltage and offset were adjusted to maximise the signal to noise ratio. Parameters were kept constant for negative control slices. Images were acquired at 600Hz and frame averaged 2-4 times, as needed. Images of patched cells were obtained with a 63X oil objective. Z-stack thickness depended on cell morphology, and was between 5-20µm. Optical slice thickness was 0.5µm. Images were visualised and processed in LAS X and FIJI. When imaging to quantify OPC proliferation or differentiation, a 20X objective was used. Z-stack thickness was determined based on OLIG2 signal (when imaging differentiation) or NG2 signal (when imaging proliferation) and was typically 25-35µm. Optical slice thickness was 1.5 or 1.6 µm. Images were analysed in Fiji and cells were counted manually. Four non-overlapping images were acquired in the cingulate cortex from two slices per animal, for a total of eight images per animal.

### Statistical analysis

Data are shown as mean±sem. Statistics were computed in GraphPad Prism or manually in Excel. When comparing two conditions, an unpaired two-tailed t-test was used; variance was tested by F-test, and Welch’s correction applied if unequal. Proportions were tested with a χ^2^ test, with Yates’ correction for small numbers. When comparing three conditions a one-way ANOVA was used; variance was tested with a Brown-Forsythe test, and Welch’s correction was applied if variance was unequal. Post-hoc comparisons were performed with a Holm-Bonferroni test.

---

**Figure S1.**
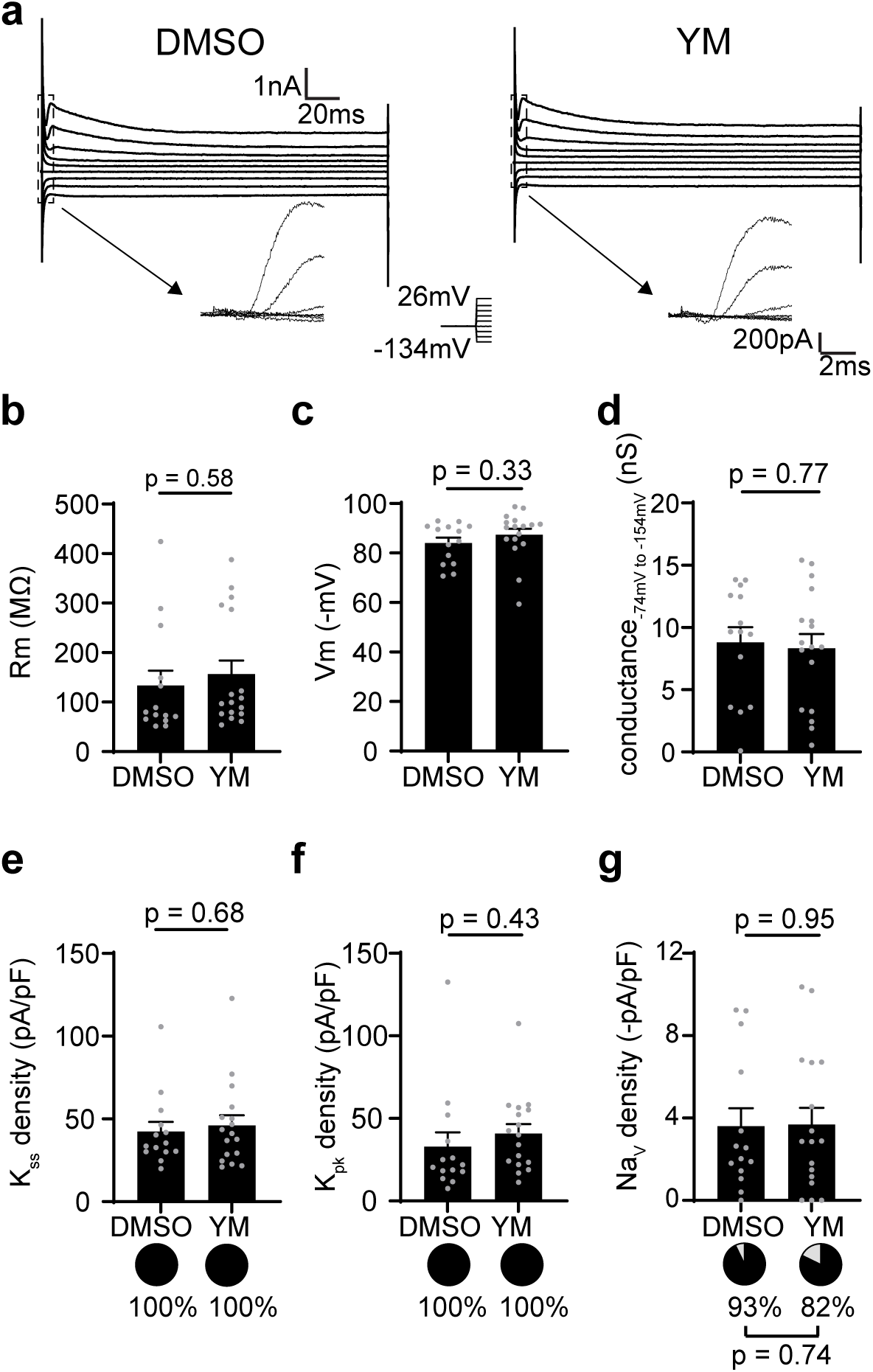
**a**, Representative current traces in response to 20 mV voltage steps between −134 mV and +40 mV in vehicle (DMSO)- and YM-treated control cortical NG2-EYFP OPCs recorded in P27-P33 mice. The arrows indicate the leak-subtracted currents during the first 7 ms of the voltage pulse. **b**-**d**, Quantification of membrane resistance (Rm, **b**), resting membrane potential (Vm, **c**), inward K^+^ conductance (conductance_−74 mV to −154 mV_, **d**) in YM-treated OPCs compared to vehicle-treated OPCs. **e**-**g**, Quantification of the density (bar graphs) and proportion (pie charts) of cells with steady-state outward K^+^ (K_ss_) currents (**e**), peak outward voltage-gated K^+^ (K_pk_) currents (**f**), and voltage-gated Na^+^ channel (Na_V_) currents (**g**) in YM-treated OPCs compared to vehicle-treated OPCs. Data are shown as mean±sem, with the grey dots indicating individual recorded cells. p-values were calculated by unpaired two-tailed t-test (bar graphs) or Yates’ corrected χ^2^ test (pie charts).

**Figure S2.**
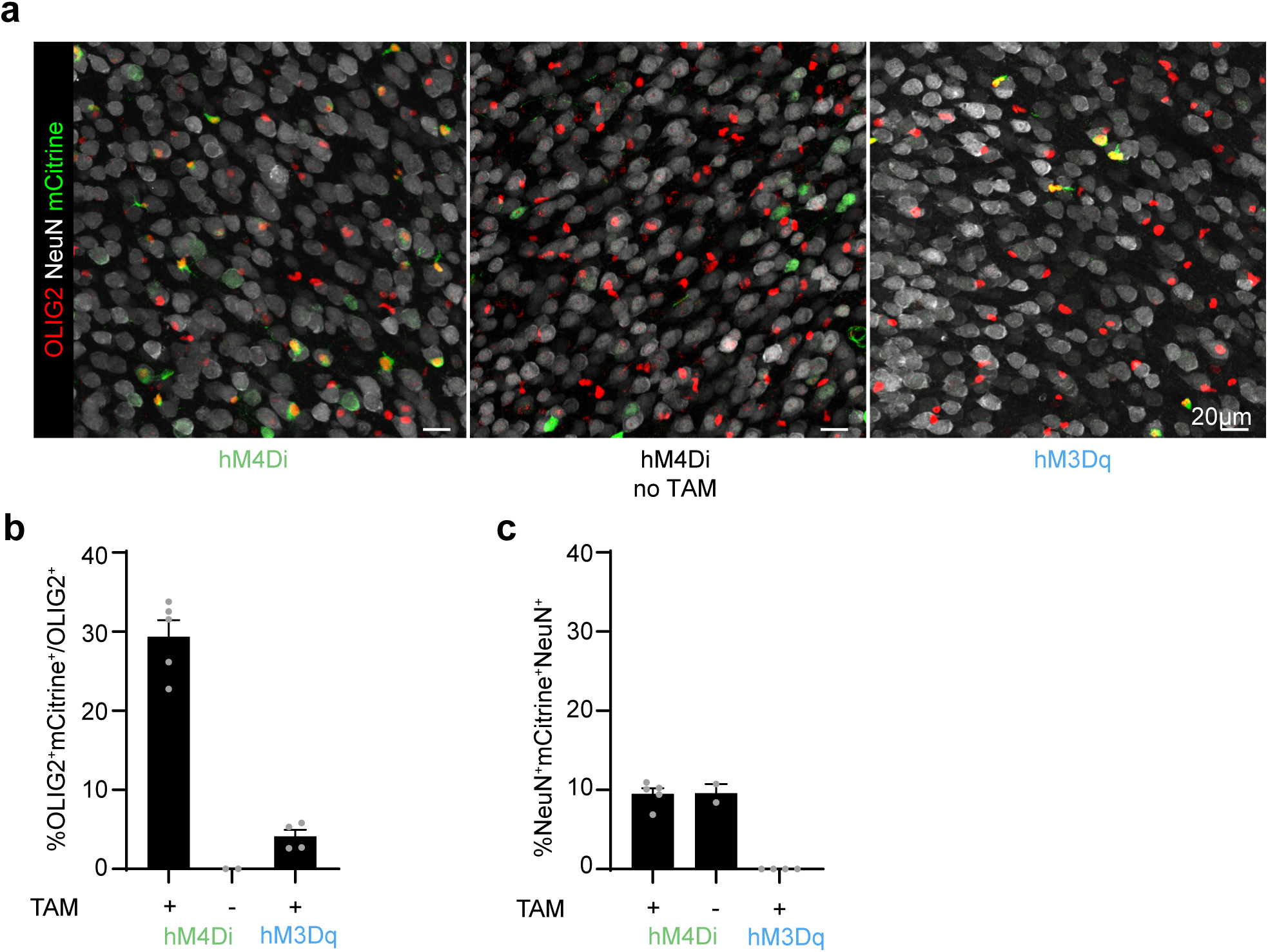
**a**, Representative images of OLIG2^+^mCitrine^+^ and NeuN^+^mCitrine^+^ cells in hM4Di (PdgfraCreER^T2^:hM4Di-mCitrine) mice given tamoxifen (left), hM4Di mice without tamoxifen administration (right) and hM3Dq (PdgfraCreER^T2^:hM3Dq-mCitrine) mice given tamoxifen. Scale bars 20 µm. **b**, Quantification of the proportion of recombined OLIG2^+^ cells in hM4Di mice dosed with tamoxifen, hM4Di mice without tamoxifen administration and hM3Dq mice given tamoxifen. **c**, Quantification of the proportion of recombined NeuN^+^ cells in hM4Di mice dosed with tamoxifen, hM4Di mice without tamoxifen administration and hM3Dq mice given tamoxifen. Data are shown as mean ±sem. Grey data points indicate individual animals.

